# Stable Vesicle-Associated BDNF from Embryonic and Young Cortical Extracellular Vesicles

**DOI:** 10.64898/2026.06.08.730839

**Authors:** Raquel García-Rodríguez, Marta Carús-Cadavieco, Irene Clares Pedrero, Carlos Cabañas, Carlos G. Dotti, Francesc X. Guix

## Abstract

In this work we show that small extracellular vesicles (sEVs) from embryonic mouse cortex or from cultured embryonic cortical neurons contain high levels of BDNF and sustain TrkB-dependent neuroprotective signaling. By contrast, sEVs from aged cortex are depleted of BDNF, and cells lacking active TrkB fail to mount a protective response when exposed to the same sEVs. Biochemical fractionation and trypsin sensitivity assay indicate that BDNF is a constitutive EV component and is exposed on or tightly associated with the vesicle surface—an arrangement that likely increases local ligand density. In a stability assay, EV-associated BDNF retained activity longer than soluble BDNF. Together, our findings suggest that many developmental effects of BDNF may be mediated by EVs, that impaired stress responses in the aged brain could reflect reduced formation of BDNF-containing EVs, and that embryonic sEVs may provide a more efficient vehicle for BDNF delivery than current therapeutic approaches.

## Introduction

Neuronal ageing is not simply a matter of cell loss, but of progressive erosion of the molecular programs that allow post-mitotic neurons to remain functional for decades. Among the numerous mechanisms that act to counteract the effects of aging [1-5], work done in the last decade has shown that one of vital importance is that played by small extracellular vesicles (sEVs). According to the current MISEV2023 recommendations, EV is the preferred umbrella term for membrane-delimited particles released by cells that do not replicate and that can transport proteins, lipids, and nucleic acids between donor and recipient cells [6]. In the nervous system, EVs are not merely by-products of membrane turnover, but active mediators of intercellular communication. Neuronal EVs influence synaptic organization, trophic signaling, and glial responses. Moreover, recent work from our laboratory has shown that biological activity of EVs is also strongly conditioned by the age and physiological state of the donor cells. Neurons aged *in vitro* increase EV secretion in association with proteostatic and endolysosomal stress, suggesting that EV release becomes part of the neuronal response to ageing-associated burden [7]. By contrast, EVs derived from young neural cultures attenuate astrocytic reactivity, supporting the concept that early-life EVs carry a trophic and homeostatic cargo distinct from that of EVs derived from older cells [8]. Together, these observations suggest that developmental stage is likely to determine whether neuronal EVs propagate stress or confer protection.

Among the pathways that sustain neuronal resilience, brain-derived neurotrophic factor (BDNF) signaling through TrkB is especially relevant because it couples trophic support to pro-survival and plasticity-related cascades, including PI3K-Akt, MAPK-Erk, and PLCγ signaling. Alterations in the BDNF/TrkB axis have been repeatedly linked to normal brain ageing, impaired synaptic plasticity, cognitive decline, and heightened susceptibility to neurodegenerative disease [9-11]. This makes the BDNF pathway an attractive mechanistic entry point for strategies aimed at preserving neuronal homeostasis in the ageing brain. Consistently, recent work has shown that sEVs can activate TrkB-dependent pathways in recipient neurons and preserve neuronal complexity under stress conditions [12, 13]. Importantly, BDNF itself can shape neuronal EV signaling: BDNF stimulation promotes the sorting of specific miRNAs into neuronal EVs, thereby enhancing circuit connectivity and synaptic maturation in recipient neurons [14].

However, despite its therapeutic appeal, the direct administration of soluble BDNF has major translational limitations. Recombinant BDNF shows poor pharmacokinetic behavior, limited tissue diffusion, restricted brain availability, and rapid loss of bioactivity in extracellular environments. These constraints have long complicated the clinical exploitation of BDNF-based therapies and have motivated the search for delivery systems that can stabilize the trophic factor while preserving its biological activity [10, 15]. In this context, any endogenous carrier able to protect BDNF from degradation and present it to recipient neurons in a biologically competent form would be of particular interest.

Building on that concept, the present study tests the hypothesis that sEVs derived from embryonic cortex or from young primary cortical cultures exert a neuroprotective effect on aged cortical neurons, and that this activity is mediated to a significant extent by vesicle-associated BDNF. The data support a model in which young-source EVs reduce neuronal damage, activate TrkB/Akt survival signaling, and preserve BDNF in a form that is substantially more stable than soluble BDNF.

## Materials and Methods

### Primary cortical cultures and *in vitro* ageing model

Primary cortical cultures were prepared from embryonic day 16 (E16) cortical tissue following the general workflow used in the source study. Briefly, cortices were dissected in ice-cold Hanks’ Balanced Salt Solution (HBSS), digested with 0.25% trypsin and 0.5 mg/mL DNase for 15 min at 37°C, washed, and mechanically dissociated in plating medium (Minimum Essential Medium supplemented with 10% horse serum and 20% glucose). The resulting cell suspension was filtered through a 70-um mesh, counted, and seeded onto poly-L-lysine-coated plates. After initial attachment, the medium was replaced with Neurobasal medium supplemented with B27 and GlutaMAX. Cultures used as aged recipient neurons were maintained until 21 days *in vitro* (DIV), whereas conditioned medium for young culture-derived EVs was collected at 7 DIV.

### Rat glioma C6 cell line cultures

Cells were cultured in Dulbecco’s Modified Eagle Medium (DMEM, Gibco) supplemented with 10% fetal bovine serum (FBS, Gibco), 100 U/mL penicillin, and 100 μg/mL streptomycin. Cultures were maintained at 37°C in a humified atmosphere containing 5% CO_2_. Cells were subcultured when they reached 80-90% confluence. For experiments, cells were seeded at a density of 1 × 10^5^ cells/mL in 6-well plates and allowed to adhere for 24 hours before treatment.

### Isolation of extracellular vesicles from embryonic cortical tissue

Small extracellular vesicles (EVs) were isolated from embryonic cortical tissue using differential centrifugation followed by sucrose-gradient enrichment, adapting the procedure described in Vella et al. [16]. Briefly, cortices were incubated in filtered Hibernate medium containing 75 U/mL type III collagenase (800 uL per 100 mg tissue) for 20 min at 37°C with intermittent inversion and gentle homogenization. The digested material was then subjected to sequential centrifugation at 300 x g for 5 min, 2,000 x g for 10 min, and 10,000 x g for 30 min at 4°C. The resulting supernatant was loaded onto a discontinuous sucrose gradient composed of 0.6 M, 1.3 M, and 2.5 M sucrose layers and ultracentrifuged at 180,000 x g for 3 h at 4°C using a swinging-bucket rotor. Individual fractions were collected, washed in filtered PBS by ultracentrifugation at 100,000 x g for 1 h, and resuspended in filtered PBS for nanoparticle tracking analysis, electron microscopy, biochemical characterization, and functional experiments.

### Isolation of extracellular vesicles from conditioned medium of young cortical cultures

EVs released by young cortical cultures were isolated from conditioned medium collected at 7 DIV. The medium was cleared by sequential centrifugation at 200 x g and 2,000 x g for 10 min each at 4°C, followed by ultracentrifugation at 10,000 x g for 30 min to remove apoptotic bodies and larger vesicles. The clarified supernatant was then layered onto a 30% sucrose cushion and ultracentrifuged at 100,000 x g for 2 h. The EV-containing fraction was recovered, washed once in sterile filtered PBS by ultracentrifugation at 100,000 x g for 1 h, and finally resuspended in sterile filtered PBS for downstream analyses.

### Transmission electron microscopy and nanoparticle tracking analysis

Purified EV preparations were characterized morphologically by transmission electron microscopy (TEM) and biophysically by nanoparticle tracking analysis (NTA). For TEM, EVs were placed on glow-discharged formvar/carbon-coated copper grids, negatively stained with 2% uranyl acetate, and imaged with a JEOL JEM-1010 transmission electron microscope. For NTA, samples were analyzed using a NanoSight NS300 instrument equipped with a 488-nm laser. Three 60-s recordings were acquired for each preparation and used to determine particle concentration and size distribution.

### EV treatments and pharmacological inhibition

For functional assays, 21 DIV cortical cultures were incubated for 24 h with embryonic cortical EVs at 1 × 10^9^ particles/mL. To evaluate the contribution of BDNF-TrkB signaling, recipient cultures were preincubated with the TrkB antagonist cyclotraxin-B (1 μM) prior to EV addition and were maintained in inhibitor-containing medium throughout the treatment interval. To assess neuroprotection under inflammatory stress, C6 glial cells were challenged with TNFα (50 ng/mL, #315-01A, PrepoTech) for 24h, both in the presence and absence of EVs (1 × 10^9^ particles/mL).

### Lactate dehydrogenase release assay

Cell damage was quantified by measuring lactate dehydrogenase (LDH) released into conditioned medium. At the end of treatment, aliquots of culture supernatant were collected and processed with the Pierce LDH Cytotoxicity Assay Kit according to the manufacturer’s instructions. Formazan production was quantified by absorbance at 490 nm using a microplate reader, and values were normalized to the appropriate control condition for each experiment.

### Western blotting

Cells and EV preparations were lysed in RIPA buffer containing protease and phosphatase inhibitors. After sonication and clarification, protein concentration was determined by BCA assay. Samples were denatured in reducing loading buffer, separated by SDS-PAGE, and transferred to nitrocellulose membranes. Membranes were blocked in BSA-containing buffer and incubated with primary antibodies directed against EV markers (including CD81, flotillin, and calnexin), survival and trophic signaling proteins (including TrkB, phospho-TrkB, and BDNF), and the corresponding loading controls. Bands were detected with HRP-conjugated secondary antibodies and quantified by densitometry in FIJI/ImageJ.

### Size-exclusion chromatography and protease accessibility of EV-associated BDNF

To determine whether BDNF was physically associated with EVs rather than representing free contaminating protein, purified vesicles from cortical cells maintained for 7 days *in vitro* (7 DIV EVs) were further fractionated by size-exclusion chromatography (SEC). Fractions were screened for EV markers and BDNF immunoreactivity by immunoblotting, and BDNF signal was interpreted in relation to the EV-rich fractions. To assess whether EV-associated BDNF was exposed at the vesicle surface, intact EV preparations were incubated with trypsin under non-permeabilizing conditions and then re-analyzed for BDNF signal and particle profile. Particle concentration and size distribution after protease treatment were re-evaluated by NTA to verify preservation of the vesicular population.

### Stability of EV-associated versus soluble BDNF

The relative stability of vesicle-associated and soluble BDNF was examined by incubating EV-bound BDNF and free soluble BDNF under matched conditions for the indicated time intervals up to 24 h. At each time point, remaining BDNF immunoreactivity was measured by immunoblotting, and signal intensity was expressed relative to the corresponding starting material. This assay was used to compare the temporal persistence of the EV-associated and free forms of the neurotrophin.

### Statistical analysis

Data are presented as mean +/-SEM. Statistical analyses were performed in GraphPad Prism. Normality was assessed with the Shapiro-Wilk test when appropriate. Two-group comparisons were analyzed using Student’s t-test, whereas multiple-group comparisons were evaluated by one-way or two-way ANOVA followed by Tukey’s post hoc test, as indicated in the figure legends. A p value < 0.05 was considered statistically significant.

## Results

### Embryonic cortical and culture-derived EV preparations show a small-EV-enriched profile

Before addressing function, EV preparations from embryonic cortex and from young primary cortical cultures were characterized morphologically and biochemically (Figure 1). EVs isolated from the cortex of embryonic day 16 mice (E16 EVs) were obtained through tissue dissociation, ultracentrifugation, and sucrose-gradient fractionation (see Materials and Methods). Electron microscopy confirmed the presence of rounded membrane-bound particles in the EV-enriched fractions, with fraction 2 (F2) being the most enriched in small-sized vesicles with diameters predominantly between 50-200 nm (Fig. 1A-B). Immunoblotting showed enrichment of EV markers such as CD81 and flotillin with absence of the endoplasmic reticulum marker calnexin in this fraction (Fig. 1C). EVs isolated from conditioned medium of young cortical cultures maintained *in vitro* for seven days (7DIV EVs) displayed a comparable size distribution by nanoparticle tracking analysis and a typical vesicular morphology by electron microscopy (Fig. 1D-E). Therefore, we next used fraction 2 of EVs isolated from embryonic brain cortex and from young cortical cultures for BDNF content analysis.

**Figure 1.**
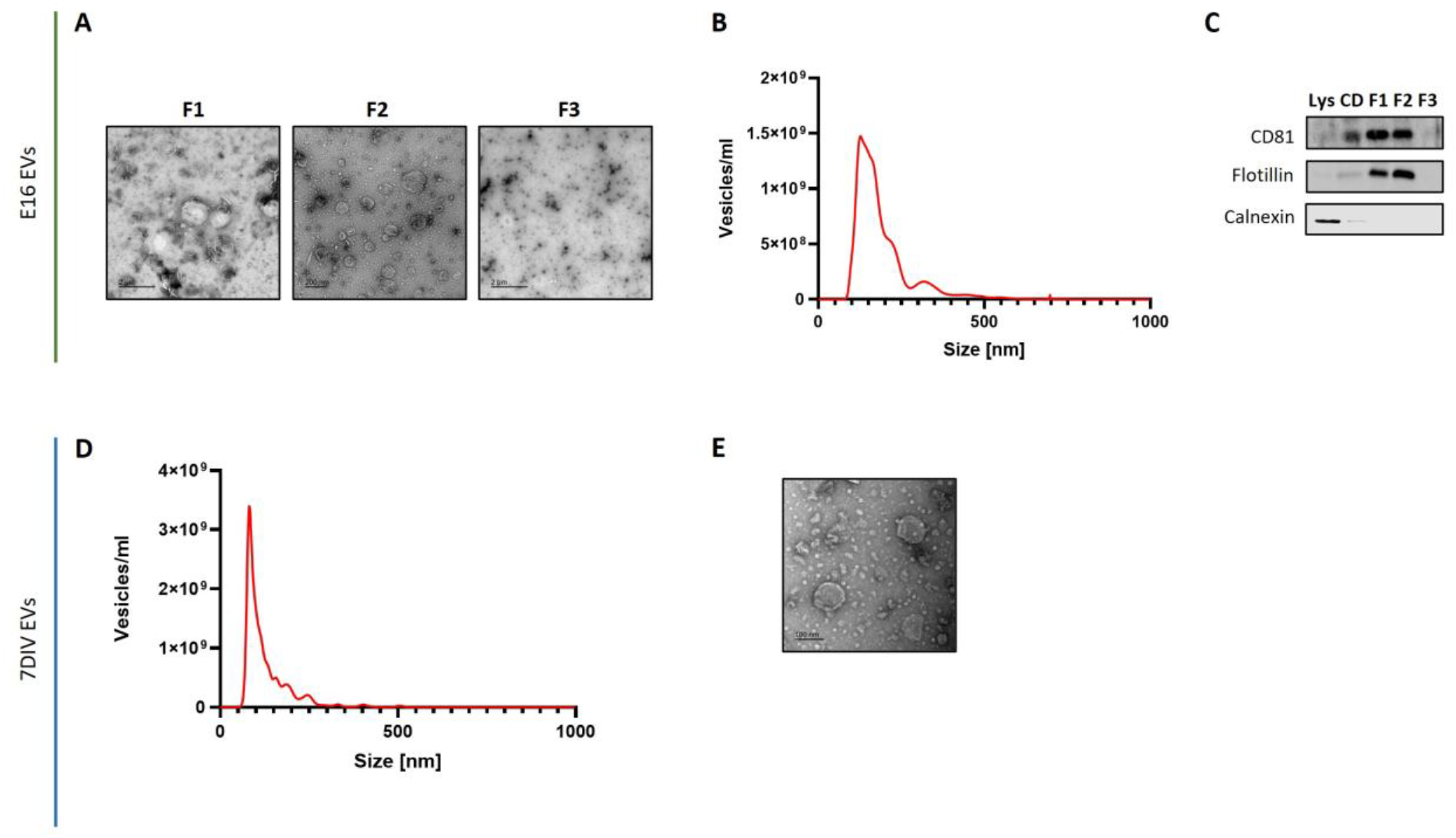
Isolation and characterization of small extracellular vesicles from mouse embryonic cerebral cortex and young neuronal cultures. (A) Representative electron microscopy images showing the different phases of the gradient used for vesicle isolation, with fraction 2 (F2) being the most enriched in small sized vesicles. (B) Representative plot showing the size distribution of the vesicles in fraction 2 (F2) analyzed by Nanoparticle Tracking Analysis (NTA). Most vesicles have a size between 50 and 200 nm. (C) Western blot analysis of the different vesicle isolation fractions, showing the enrichment of EV-specific markers such as CD81 and flotillin, and the absence of the endoplasmic reticulum marker calnexin in the fraction enriched in small vesicles (F2). (D) Representative size distribution of the vesicles present in the conditioned medium of mouse neuronal cultures and analyzed by Nanoparticle Tracking Analysis (NTA). (E) Representative transmission electron microscopy (TEM) image of vesicles obtained from the medium of mouse neuronal cultures by centrifugation at 100,000 x g. Lys=lysate; CD=cell debris.

### Embryonic sEVs have significantly more BDNF than sEVs from older brains: contribution to neuronal survival

Given the prominent role of BDNF in neuronal survival and plasticity [17-21], the trophic content of EV preparations was examined next. Immunoblotting revealed that EVs isolated from young cortical cultures (7DIV EVs) are enriched in mature BDNF and lacked detectable proBDNF, while embryonic cortical EVs (E16 EVs) also contained clear mature BDNF signal together with EV markers (Fig. 2A). In contrast, EVs derived from aged 20-month-old cortex (20 m EVs) contained substantially lower levels of BDNF than embryonic EVs (Fig. 2B).

**Figure 2.**
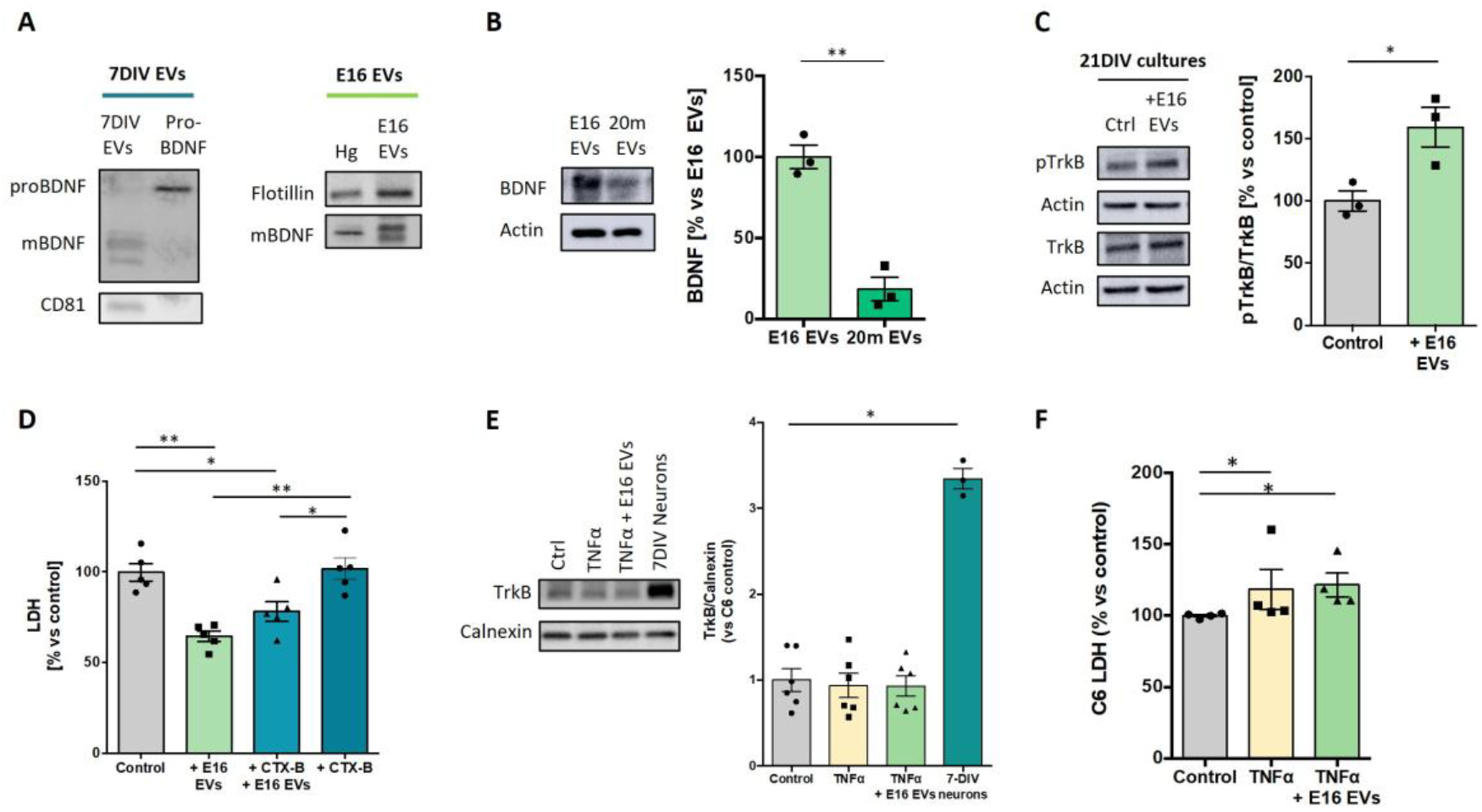
BDNF is detected in EVs from young cortical cultures and embryonic cortex, is markedly reduced in EVs isolated form aged cortex and triggers TrkB activation in aged cortical neurons. (A) Western blot confirming the presence of mature BDNF in both purified vesicles from cortical cells maintained for 7 days in vitro (7DIV EVs) and cortical cells from 16-day-old embryos (E16 EVs). The absence of the precursor form, pro-BDNF, is also shown in 7DIV EVs. Hg=homogenate (B) Western blot analysis showing lower levels of mature BDNF in purified vesicles from 20-month-old mice (20m EVs) compared with E16 EVs. Actin was used as loading control. Data represent the mean **±** SEM as a percentage of E16 EV values. n = 3; *p<0.05; Student’s t-test. (C) Western blot analysis of the ratio of activated TrkB (pTrkB) to total TrkB protein in 21DIV cortical cultures treated or not with E16 EVs (n = 3; *p<0.05; Student’s t-test). Actin was used as a loading control. (D) Absorbance measurement of LDH in conditioned medium of 21DIV cortical cultures treated or not with cyclotraxin B (CTX-B, a selective TrkB inhibitor) and E16 EVs (n = 5; *p < 0.05 and **p < 0.01; two-way ANOVA followed by Tukey’s post hoc correction). Note that the addition of the TrkB inhibitor alone had no effect and only partially prevents the effect of the EVs. (E) Western blot analysis of total TrkB protein in C6 glial cells treated or not with TNFα (50 ng/mL) in the presence or absence of E16 EVs. TrkB levels in 7 DIV neurons were also compared with those in C6 cells. Calnexin was used as a loading control (n = 6; *p < 0.05; one-way ANOVA followed by Tukey’s post hoc correction). (F) Absorbance measurement of LDH release into the conditioned medium of C6 glial cells treated or not with TNFα (50 ng/mL) in the presence or absence of E16 EVs (n = 4; *p < 0.05; one-way ANOVA followed by Tukey’s post hoc correction). All data represent the mean **±** SEM as a percentage of the control (untreated) values.

To test whether the trophic content of embryonic EVs engages canonical BDNF signaling, TrkB activation was analyzed in recipient 21 DIV cortical cultures. Treatment with E16 EVs increased the ratio of phosphorylated TrkB to total TrkB, demonstrating activation of the receptor pathway (Fig. 2C). Moreover, EVs incubation reduced LDH levels in the culture medium (Fig. 2D), suggesting a neuroprotective role. In order to establish the possible causal relationship with the vesicles’ BDNF, 21 DIV cortical cultures were co-incubated with EVs and cyclotraxin-B, a selective TrkB inhibitor [22]. This treatment attenuated the EV-induced reduction in LDH release, whereas cyclotraxin-B alone did not affect basal LDH levels (Fig. 2D). These data indicate that TrkB signaling triggered by E16 EVs is not merely a passive consequence of EV treatment but requires the presence of BDNF on the vesicles. To further validate this mechanism, we utilized C6 glial cells, which primarily express the truncated TrkB.T1 isoform-lacking the intracellular kinase domain-rather than full-length pro-survival TrkB receptor [23]. In these cells, which exhibit reduced full-length TrkB expression compared to primary cortical neurons (Fig. 2E), E16 EVs failed to mitigate TNF-α-induced LDH release (Fig. 2F). While these results identify BDNF as a key functional component, the incomplete loss of protection after TrkB inhibition suggests that EV-mediated neuroprotection is multifactorial and that BDNF/TrkB represents an important, but not exclusive, effector arm.

### EV-associated BDNF co-fractionates with EVs, is surface-accessible, and is more stable than soluble BDNF

The physical relationship between BDNF and EVs was then examined in more detail. Following size-exclusion chromatography of EVs derived from cortical cells maintained *in vitro* for 7 days (7DIV EVs), BDNF signal co-eluted with CD81-positive EV-rich fractions, arguing against the possibility that the detected trophic factor merely reflects contaminating free protein (Fig. 3A). Protease digestion experiments further showed that trypsin treatment abolished the EV-associated BDNF signal (Fig. 3B) without substantially altering vesicle integrity, particle number or size distribution (Fig. 3C). This pattern is consistent with BDNF being exposed at, or tightly associated with, the external EV surface rather than being protected exclusively within the vesicle lumen. Finally, a direct comparison of vesicle-associated and soluble BDNF demonstrated a marked stability advantage for the EV-associated form: BDNF linked to EVs retained most of its signal over 24 hours, whereas soluble BDNF declined rapidly over the same period (Fig. 3D). These findings provide a mechanistic explanation for how young-source EVs may amplify trophic efficacy: they do not simply carry BDNF but preserve it in a configuration with substantially prolonged extracellular persistence.

**Figure 3.**
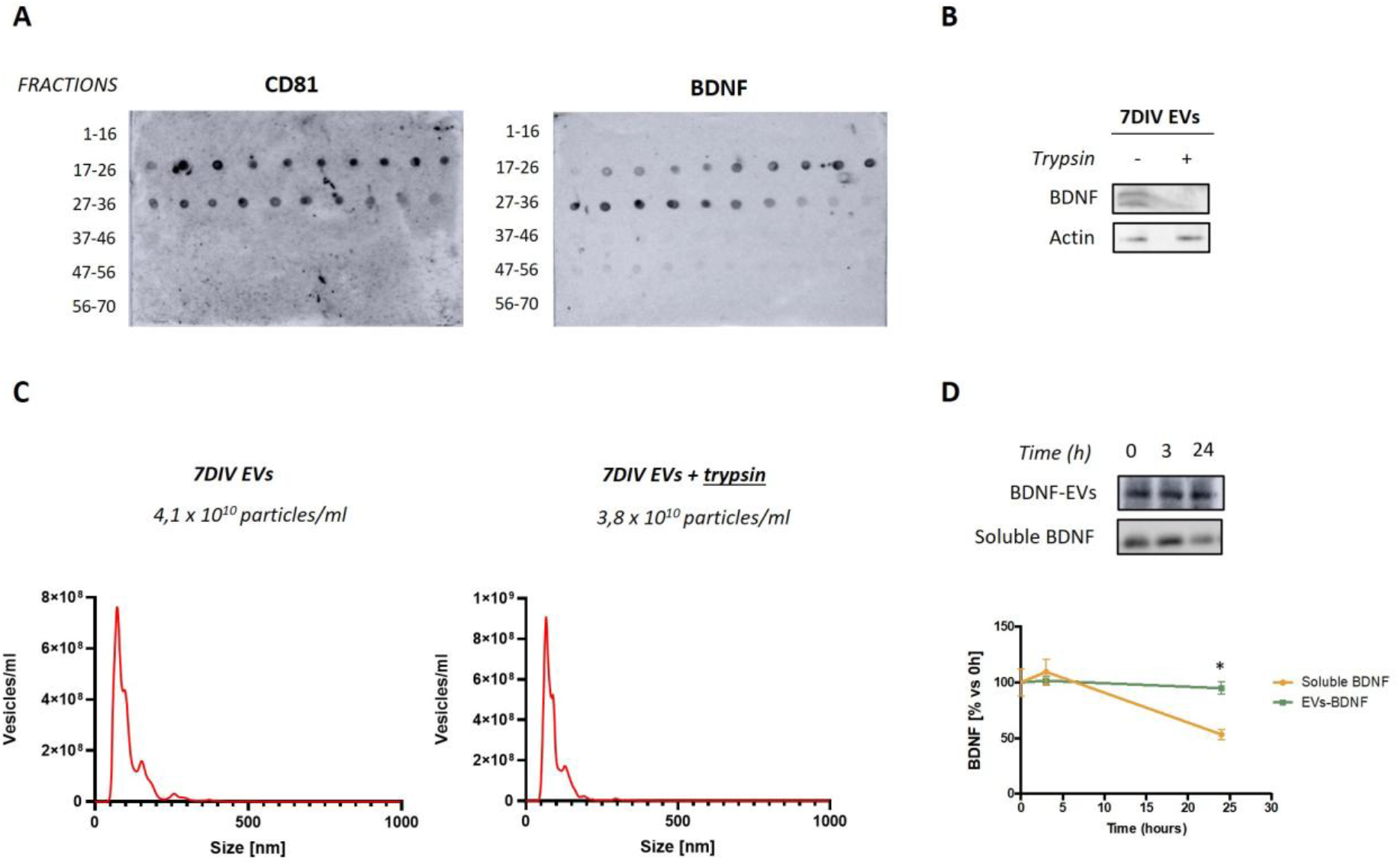
BDNF in embryonic EVs derived from donor cells is present on the outer surface and exhibits remarkable stability. (A) Size exclusion chromatography of extracellular vesicles derived from cortical cells maintained in vitro for 7 days (7DIV EVs). Fractions were analyzed by dot blot and revealed the presence of BDNF exclusively in EV-rich fractions, as confirmed by the expression of the specific marker CD81. (B) Western blot analysis of the mature form of BDNF in 7DIV EVs, showing its loss after treatment of the EVs with 45 μg/mL trypsin for 30 min at 37°C. (C) Particle size distribution and concentration measured by NTA before and after trypsin treatment. (D) Western blot analysis of the stability of BDNF present in embryonic EVs (E16 EVs) compared with soluble BDNF. The presence of the mature form of BDNF was assessed at three different time points: 0, 3, and 24 h. Data show the mean **±** SEM expressed as the percentage of BDNF levels normalized to the values measured at 0 hours (n = 3; *p < 0.05; one-way ANOVA followed by Tukey’s post hoc correction)

## Discussion

The present work identifies BDNF in extracellular vesicles released by embryonic cortex or by young primary cortical neurons as a trophic system capable of preserving the viability of cortical neurons in a stressful environment. Across the figure set, the protective phenotype is internally consistent: young-source EVs contain substantially more BDNF than EVs from old cortex, activate TrkB in recipient neurons, and retain BDNF signal longer than the free soluble protein. These data, beyond reinforcing the notion that BDNF from embryonic stages has an important role in cell survival during brain development, demonstrate that the effect would require BDNF to be embedded in sEVs. As a matter of fact, the partial loss of protection after TrkB blockade places the BDNF/TrkB axis in a mechanistically relevant position rather than as a simple correlate of EV exposure.

In other systems, exosome-associated BDNF has been implicated in protection against TNFα-induced neuronal apoptosis, diabetic retinal neurodegeneration, ischemic injury, Parkinsonian degeneration, and spinal cord trauma [24-28]. Most of those studies relied on EVs isolated from glial, mesenchymal stem cell, or engineered EV sources. By contrast, the present data indicate that an endogenous neural developmental source—embryonic cortex or young cortical neurons—already contains a naturally assembled trophic EV program able to engage aged neurons. This distinction is important conceptually, because it suggests that the beneficial signal is not necessarily an artificial product of engineering, but may reflect a physiological mechanism by which young neural tissue distributes survival/differentiation cues to neighboring cells.

A particularly informative aspect of the present dataset is the topology and stability of vesicle-associated BDNF. The co-elution of BDNF with EV-rich fractions and the loss of signal after trypsin treatment are most consistent with BDNF being exposed on, or tightly associated with, the outer vesicle surface. Such an arrangement could favor efficient receptor engagement by increasing local ligand density and allowing immediate access to TrkB upon vesicle-cell contact. At the same time, EV association appears to prolong extracellular persistence of the trophic factor, providing an attractive mechanistic explanation for why vesicle-bound BDNF may outperform free BDNF in biological settings where soluble protein is rapidly diluted, degraded, or otherwise inactivated [10, 15, 27, 28]. This interpretation is also relevant for therapeutic design, because it suggests that the vesicle membrane is not merely a passive carrier but part of the mechanism that preserves ligand competence.

The age dependence of the effect is equally notable. EVs from old cortex were markedly depleted of BDNF, consistent with the idea that neuronal ageing changes EV composition and function, potentially through age-related alterations in membrane fluidity and stability [29], which may impair BDNF retention on the EV surface. Previous work showed that neurons aged *in vitro* increase EV secretion in association with endolysosomal and cholesterol-related stress, whereas EVs from young neural cultures can dampen adverse glial responses [7, 8]. Our findings add a trophic dimension to that framework by indicating that ageing is accompanied not only by changes in EV quantity, but also by loss of a survival-promoting EV-associated neurotrophin signal. An additional point of interest is that not all EV-associated BDNF signals are expected to be beneficial: EVs enriched in proBDNF have been reported to promote neuronal apoptosis in other contexts, emphasizing that the biological outcome likely depends on the maturation state, presentation mode, and cellular origin of the neurotrophin cargo [30].

Several limitations should be considered. First, the present figures support a central role for EV-associated BDNF, but they do not fully exclude contributions from other vesicular proteins, lipids, or RNAs; indeed, the incomplete suppression produced by cyclotraxin-B argues that the protective program is probably multifactorial. Second, the data do not yet resolve the relative contribution of mature BDNF versus proBDNF within the active EV preparations, nor do they establish whether vesicle-bound BDNF is transferred to the recipient membrane or acts exclusively from the EV surface. Third, although the stability assay strongly suggests a pharmacological advantage for EV-associated BDNF, this was assessed *in vitro* and should be extended to more physiological extracellular environments and, ultimately, *in vivo*. Future work should therefore combine BDNF neutralization or immunodepletion strategies, quantitative proteomics, and *in vivo* testing in brain ageing or neurodegeneration models to define how far young neural EVs can be developed as a biologically grounded neuroprotective platform.

Overall, the data support the view that young neural EVs function as vehicles for stabilized trophic signaling and that BDNF is a major component of this effect. The observation that EV association preserves BDNF and couples it to TrkB-dependent protection in aged neurons provides a mechanistic bridge between developmental neurobiology, extracellular vesicle biology, and brain ageing research. More broadly, these results raise the possibility that part of the rejuvenating influence of young neural environments may be mediated by vesicular neurotrophin presentation. Defining the molecular architecture of this trophic EV signal should therefore be of considerable interest both for understanding neuronal ageing and for designing next-generation neuroprotective interventions.

## Conclusions

Across the figure set, the data support a coherent model in which EVs produced by embryonic cortex or young cortical cultures act as trophic conveyors, whereas EVs from aged cortex are comparatively deficient in this activity. The protective response in aged recipient neurons tracks with activation of TrkB, enrichment of EV-associated BDNF, and increased stability of the vesicle-bound trophic factor. Together, these results position vesicle-associated BDNF as a central mechanistic component of the neuroprotective activity of young-source cortical EVs.

Across the figure set, the data define a coherent model: EVs from embryonic or young cortical tissue function as potent trophic conveyors, whereas EVs from aged cortex are markedly deficient. The durable protective response in aged recipient neurons correlates with TrkB activation, enrichment of EV-associated BDNF, and enhanced stability of vesicle-bound BDNF, highlighting translational potential.

## AUTHORS’ CONTRIBUTION

R.G-R. performed the majority of the experiments, collected data and conducted the statistical analysis; M.C-C. performed experiments using the C6 cell line; I.C.P. and C.C. advised on and contributed to the isolation of EVs by size-exclusion chromatography; C.G.D. and F.X.G. designed the study and conceptualized the research; F.G.X wrote the initial draft; C.G.D, F.G.X and R.G-R. reviewed and edited the manuscript. All authors have read and approved the final manuscript.

## FUNDING

This work was supported by Agencia Estatal de Investigación (MICIU/AEI/10.13039/501100011033) through grants PID2019-104389RB-I00 (ERDF/EU) to C.G.D, including the award of a predoctoral fellowship to R.G-R. for the training of research personnel (FPI) and PID2022-138334OB-I00 (ERDF/EU) to C.G.D and F.G.R.

## ACKNOWLEDGMENTS

We thank the different scientific services of the Centro de Biología Molecular Severo Ochoa (CBM): the Electron Microscopy Service for sample staining and processing and the Animal Facility for the care of the experimental animals.

## DATA AVAILABILITY

All data generated or analysed during this study are included in this published article

## ETHICS APPROVAL AND CONSENT TO PARTICIPATE

Mice were housed at the Centro de Biología Molecular “Severo Ochoa” (CSIC). All experiments were performed in accordance with European Union guidelines (2010/63/UE) regarding the use of laboratory animals.

## CONSENT FOR PUBLICATION

Not applicable.

## COMPETING INTERESTS

The authors declare no competing interests.

## Notes

### Competing Interest Statement

The authors have declared no competing interest.

